# Effects of tenascin-C on dental pulp tissue in mice *in vivo* and on the proliferation and differentiation of dental pulp stem cells into odontoblasts and calcification *in vitro*

**DOI:** 10.1101/2024.02.19.580638

**Authors:** Kentaro Kojima, Yoshihiko Akashi, Kei Nakajima, Katsutoshi Kokubun, Seikou Shintani, Kenichi Matsuzaka

**Affiliations:** Department of Pediatric Dentistry, Tokyo Dental College, Tokyo, Japan; Department of Pathology, Tokyo Dental College, Tokyo, Japan

## Abstract

This study aimed to investigate the effects of tenascin-C (TN-C) on dental pulp tissue and on dental pulp stem cells (DPSCs). In in vivo studies, A collagen sponge with phosphate-buffered saline (PBS) for the control group or with TN-C for the experimental group was placed over the dental pulp of mice. The root pulp was excised at 7 and 21 days postoperatively and was observed microscopically using HE staining and immunohistochemistry. Inflammatory cells were found in the entire pulp tissue in the control group but no inflammatory cells were identified in the pulp tissue in the TN-C-treated experimental group. Further, nestin-positive cells at 7 days and dentin sialophosphoprotein (DSPP)-positive cells at 21 days were seen in the experimental group. In in vitro studies, DPSCs were cultured in a medium with or without TN-C, after which the proliferation rate of DPSCs was measured mRNA expression levels were examined using quantitative reverse transcription polymerase chain reaction (qRT-PCR), and the formation of calcified nodules was investigated using alizarin red staining. The cell proliferation rate was not significantly different between the experimental and control groups. The expression of nestin mRNA on day 7 was significantly higher in the experimental group than in the control group (P<0.05), but the expression of osteocalcin (OCN) mRNA was significantly higher in the control group than in the experimental group (P<0.05). More calcified nodules formed in the control group than in the experimental group. These results suggest that TN-C regulates inflammation during the healing process in the dental pulp and induces the differentiation of dental pulp into odontoblast-like cells. Further, TN-C promotes the early differentiation of DPSCs into odontoblast-like cells, which suggests that TN-C may further contribute to the inhibition of excessive dentin formation.

## INTRODUCTION

The dental pulp is a loose connective tissue isolated from the external environment by enamel and dentin, that has a vascular system, nerves and a mixed population of cells including fibroblasts, odontoblasts and dental pulp stem cells (DPSCs) [1, 2, 3]. DPSCs are mesenchymal stem cell marker-positive cells that play an important role in reparative dentinogenesis and in dentin bridge formation [4, 5, 6]. Stem cells have two distinct abilities: the ability for self-renewal and the ability to differentiate into any type of cell in an organism [7, 8]. Regenerative medicine using stem cells is currently attracting increased attention [9, 10], but there are many challenges in the clinical applications of totipotent and pluripotent stem cells [11, 12, 13]. DPSCs are attracting special attention for their potential applications in regenerative medicine due to their easy accessibility and broad capacity for differentiation [14–17]. Currently, several treatment modalities exist that take advantage of the ability of DPSCs to differentiate into odontoblasts, including direct pulp capping, pulpotomy and indirect pulp capping [18, 19, 20]. In addition, regenerative endodontics has recently been considered a new treatment for pulp inflammation and necrosis in immature teeth [21, 22]. Many research studies in recent years have focused on investigating the use of growth factors such as bone morphogenetic protein (BMP), vascular endothelial growth factor (VEGF) and platelet-derived growth factor (PDGF) to promote the differentiation of DPSCs more efficiently, and the extracellular matrix has attracted attention as an important component [23, 24, 25].

Tenascin-C (TN-C) is an extracellular matrix protein with epidermal growth factor-like domains, a string of fibronectin-type III (FNIII) repeats and a fibrinogen-like globe [26, 27]. TN-C is expressed during tissue morphogenesis and wound healing and has been attracting attention as an extracellular matrix component with various functions that impact cell proliferation and differentiation [28]. Accordingly, TN-C is expected to have applications in regenerative medicine [29, 30]. Furthermore, TN-C has been reported to be expressed during dentin formation in tooth development and during tertiary dentinogenesis following direct pulp capping [31, 32, 33]. TN-C promotes the proliferation and the differentiation of dental pulp cells to mineralized tissue-forming cells [34]. Thus, TN-C may play an important role in the differentiation of DPSCs to odontoblasts and we thought that the treatment of DPSCs with TN-C would promote odontoblast differentiation and would help improve future treatment outcomes.

Therefore, for the *in vivo* study, the root pulp was placed in phosphate-buffered saline (PBS) with or without TN-C and postoperative morphology was observed using a light microscope. For the *in vitro* study, hDPSCs were cultured with or without TN-C, after which the proliferation rate of DPSCs was measured. mRNA expression levels of an undifferentiated marker (POU5F1), an early (NESTIN) and a late (DSPP) differentiation marker of odontoblasts and a calcification marker (OCN) were examined using quantitative reverse transcription polymerase chain reaction (qRT-PCR). The formation of calcified nodules was investigated using alizarin red staining. Thus, the purpose of this study was to evaluate the effects of TN-C on dental pulp tissue and on DPSCs.

## MATERIALS AND METHODS

### in vivo study

#### Animals

The manuscript of this animal study has been written according to Preferred Reporting Items for Animal studies in Endodontology (PRIASE) 2021 guidelines. This study was conducted in compliance with the Guidelines for the Treatment of Experimental Animals (School of Dentistry, Approval Number: 220501). C57BL/6 female mice (7-week-old, each weighing approximately 20 g) were used in this study. No animals suffered infection or death during the experimental period.

#### Pulp chamber procedures

Cavities were produced in the maxillary first molars of mice, under general anesthesia with a mixture of medetomidine hydrochloride (0.75 mg/kg), midazolam (4 mg/kg), and butorphanol tartrate (5 mg/kg), using a carbide bur (M1/4; Mani, Inc., Tochigi, Japan). The coronal pulp was then excavated completely with a carbide bur, a steel round bur (ST1 CA 006; Hager and Meisinger GmbH, Berlin, Germany) and a micro-excavator (NEW OK Micro Exca 25 ° 21 mm 0.5 mm; Sedo Mfg. Co., Ltd, Ibaraki, Japan). The TN-C solution was prepared by diluting TN-C to 300 ng/mL in PBS (Gibco, city, state, USA) for the experimental group (n=3), with PBS only used as the control group (n=3), and was soaked in finely chopped Atelocollagen, Honeycomb Disc 96 (CSH-96, KOKEN, Tokyo, Japan), and placed over each pulp stump. Glass ionomer cement (Ionosit-Baseliner; DMG, Hamburg, Germany) was used to seal each defect.

#### Histological and immunohistochemical observations

At 7 and 21 days after the surgery, the mice in each group were euthanized by cervical dislocation after general anesthesia with a mixture of medetomidine hydrochloride (0.75 mg/kg), midazolam (4 mg/kg), and butorphanol tartrate (5 mg/kg) or by inhalation with isoflurane. Their entire maxilla were removed and immersed in 4% paraformaldehyde for 1 day. Each maxilla was then washed with PBS and demineralized with 10% ethylenediaminetetraacetic acid (EDTA) at 4°C for 2–3 weeks. After demineralization, the specimens were embedded in paraffin and cut into 4 μm sagittal sections using a microtome. The sections were stained with hematoxylin and eosin and were observed with a light microscope. Immunohistochemical staining was performed as follows: Sections were immersed in methanol containing 0.3% hydrogen peroxide at room temperature to block endogenous peroxidase activity, and then were blocked for 60 min with 10% goat serum at room temperature to reduce non-specific binding. Subsequently, a mouse anti-nestin monoclonal antibody (diluted 1:200; ab6142; Abcam Inc., Cambridge, UK) was used at 7 days postoperatively, and a mouse anti-DSPP monoclonal antibody (diluted 1:50; sc73632; Santa Cruz Biotechnology Inc., Santa Cruz, CA, USA) was used at 21 days postoperatively as early and late markers of odontoblast differentiation, respectively. MACH 2 Universal HRP Polymer Detection (BRR522G, BIOCARE MEDICAL, city, CA, USA) was used as the peroxidase-conjugated secondary antibody. After that, the sections were stained with DAB and observed using a light microscope. As a negative control, 1% goat serum was used instead of the primary antibody.

### in vitro study

#### Cells

hDPSCs (PT-5025; Lonza, Basel, Switzerland) derived from human dental pulp tissues were purchased from Lonza for use in this study. Those cells expressed CD29, CD73, CD90, CD105 and CD166 as surface markers of mesenchymal stem cells, and were negative for CD34, CD45 and CD133 as expected. Those cells were cultured in DPSC growth medium (DPSC-GM) by adding the contents of a DPSC SingleQuots Kit (PT-4516; Lonza) to DPSC Basal Medium (DPSC-BM) (PT-3927; Lonza) in a humidified atmosphere (37°C and 5% CO_2_). Cells at passages 8-9 (P8-9) were used in this study.

#### Preparation of TN-C in cell culture medium

Recombinant human TN-C (50 μg) (3358-TC-050; R&D Systems Inc, Minneapolis, MN, USA) was dissolved in 100 μL PBS to produce a stock solution (500 μg/mL). The cell culture medium of the experimental group was diluted to a final concentration of 300 ng/mL TN-C by adding the above stock solution to DPSC-GM.

#### Cell proliferation assay

hDPSCs were seeded in 24-well plates at a density of 10,000 cells/cm^2^ and were cultured in DPSC-GM as a control group and in DPSC-GM + TN-C at a concentration of 300 ng/mL as the experimental group in a humidified atmosphere (37°C and 5% CO_2_). Cell proliferation was evaluated using a Cell Counting Kit-8 (Dojindo, Kumamoto, Japan). After 1, 3 and 5 days of culture, 50 μL of Cell Counting Kit-8 was added to each well and was incubated at 37°C and 5% CO_2_ for 1 hour. The resulting supernatant was aliquoted at 100 µl into each well of a 96-well plate, and absorbance was measured at 450 nm using a microplate reader (Bio Tek Gen5, Bio Tek, city, state, USA) (n=4). The results are expressed as the ratio of absorbance value relative to the baseline at day 1.

#### Odontogenic differentiation

For the induction of odontoblastic differentiation in hDPSCs, an odontogenic medium was prepared by adding 50 μg/mL L-ascorbic acid phosphate, 100 nmol/L dexamethasone, 10 mmol/L β-glycerol phosphate, 15 ng/mL BMP-4 (314-BP-050/CF; R&D Systems Inc) and 125 ng/mL Fibroblast growth factor (FGF) -8b (423-F8-025/CF; R&D Systems Inc) to DPSC-BM as previously reported [35]. hDPSCs were seeded at a density of 10,000 cells/cm^2^ and were cultured in DPSC-BM for 1 day to establish the cells and then changed to odontogenic medium the next day, after which the odontogenic medium was changed twice a week.

#### Quantitative reverse transcription polymerase chain reaction (qRT-PCR)

On days 7 and 21 of culture, hDPSCs from each group were collected. Total RNA was extracted from each sample using a RNeasy Mini Kit (QIAGEN, Hilden, Germany), after which complementary DNA (cDNA) was reversed transcribed using ReverTra Ace qPCR RT Master Mix with gDNA Remover (TOYOBO, Osaka, Japan). qRT-PCR was then performed using TaqMan Gene Expression Assays (Applied Biosystems, Foster City, CA, USA) (n=4). The following target genes were used in this study. POU Class 5 Homeobox 1 (*POU5F1*) (Hs04260367_gH) as an undifferentiated marker, *nestin* (Hs04187831_g1) as an early differentiation marker of odontoblasts, dentin sialophosphoprotein (*DSPP*) (Hs00171962_m1) as a late differentiation marker of odontoblasts, and osteocalcin (*OCN*) (Hs01587814_g1) as a calcification marker. Glyceraldehyde-3-phosphate dehydrogenase (*GAPDH*) (Hs03929097_g1) was used as an endogenous control. qRT-PCR was performed using a 7500 Fast Real-Time PCR System (Applied Biosystems). *GAPDH* mRNA expression levels were used as a baseline and target gene expression levels were corrected for relative quantitative analysis.

#### Alizarin Red Staining

After culturing hDPSCs in odontogenic medium, control and experimental groups were evaluated for calcium production at days 14 and 21 by alizarin red staining (n=4) as follows: cells were washed twice with PBS, then were fixed with 4% paraformaldehyde for 10 minutes, after which they were washed twice with Milli-Q Water and stained with Alizarin Red S solution (pH 6.3) for 10 minutes. Finally, cells were washed three times with Milli-Q Water. After staining, the ratio of the stained area was imaged using ImageJ FIJI software program (National Institutes of Health, Bethesda, MD, USA).

#### Statistical Analysis

Data were compared between the experimental and control groups using the Mann–Whitney U test. Data are presented as means ± standard deviation. A value of *P* < 0.05 is considered to be statistically significant.

## RESULTS

### in vivo study

#### Histological observations

On postoperative day 7, the odontoblast layer was disordered, and inflammatory cells such as neutrophils and macrophages were observed in the pulp tissue in the control group (Fig 1a). On the other hand, granulation tissue was seen in the dental pulp just below the pulp stump, and a vascular extension was observed but no inflammatory cell infiltration was identified in the pulp tissue in the TN-C-treated experimental group (Fig 1b). On postoperative day 21, inflammatory cell infiltration was observed in the pulp tissue within the root canal in the control group (Fig 1c). In contrast, inflammatory cell infiltration was not observed in the pulp tissue within the root canal and spindle-shaped cells were observed at the root canal opening in the TN-C-treated experimental group (Fig 1d).

**FIGURE 1.**
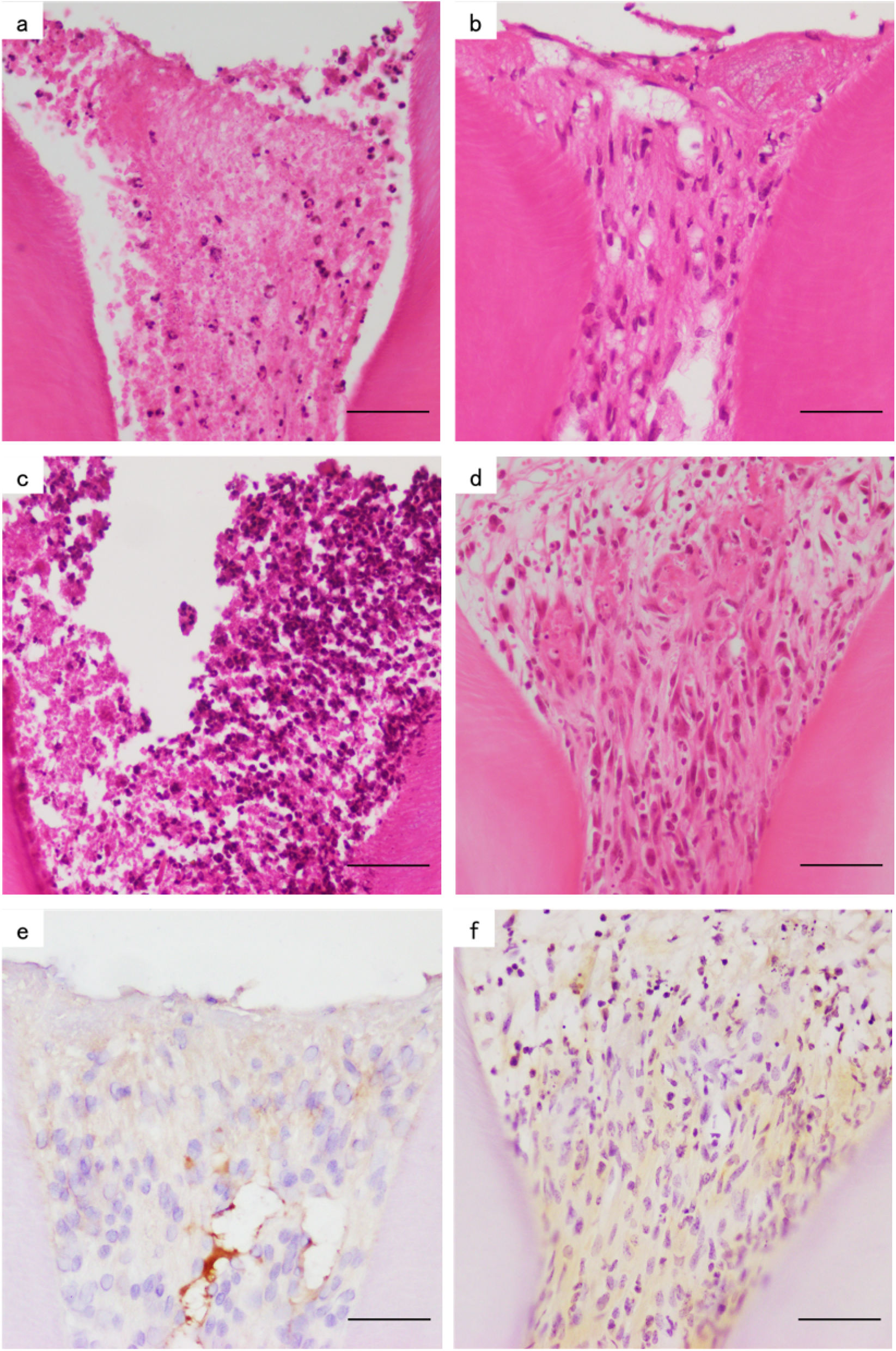
Morphological findings *in vivo*.

#### Immunohistochemistry

Nestin-positive cells were observed in the pulp just below the TN-C-containing atelocollagen contact area in the experimental group at postoperative day 7 (Fig 1e). Further, DSPP-positive cells were observed in the entire dental pulp tissue in the experimental group at postoperative day 21 (Fig 1f).

On day 7, the odontoblast layer was disordered and inflammatory cells including neutrophils and macrophages were observed in the pulp tissue in the control group (a). The pulp in the TN-C-treated experimental group contained granulation tissue in the dental pulp just below the pulp stump and vascular extensions were observed but there were no inflammatory cells (b). On day 21, an inflammatory cell infiltration was observed in the entire pulp tissue within the root canal in the control group (c). The pulp in the TN-C-treated experimental group was observed to contain spindle-shaped cells at the root canal opening but no inflammatory cells (d). On day 7, nestin-positive cells were observed in the pulp just below the TN-C-containing atelocollagen contact area in the experimental group (e). On day 21, DSPP-positive cells were observed in the experimental group (f). (Scale bars: 50 μm)

### in vitro study

#### Cell proliferation assay

With the proliferation rate at day 1 used as the baseline value, there was no significant difference in the proliferation rates of DPSCs on days 3 or 5 between the TN-C-treated experimental group and the control group (Fig 2).

**FIGURE 2.**
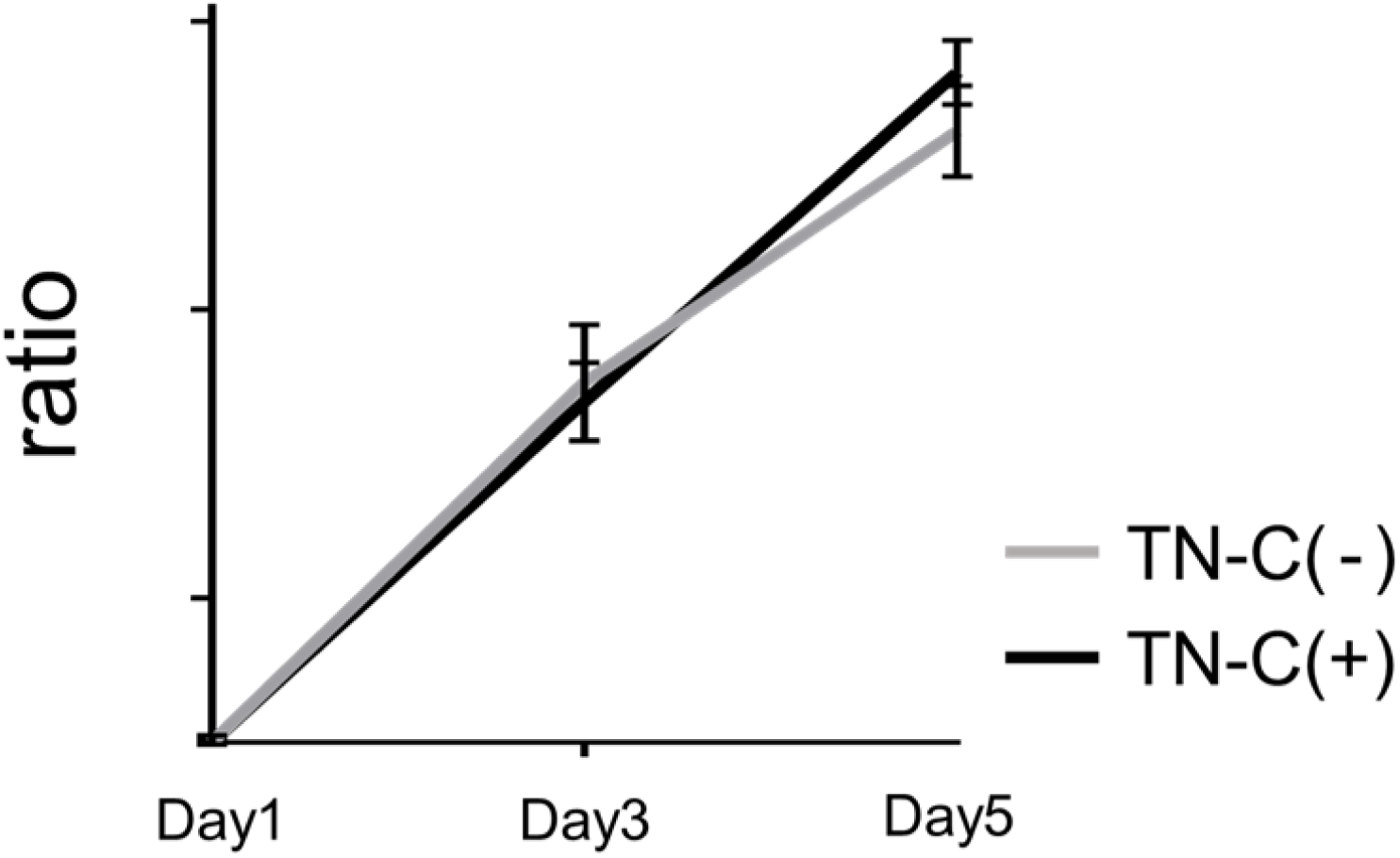
Proliferation rates of DPSCs *in vitro*.

Proliferation rates are shown relative to the baseline value (day 1) in the experimental and control groups. No significant difference was observed in cell proliferation rates between the experimental and control groups at days 3 or 5.

#### qRT-PCR analysis

The expression of *POU5F1* mRNA was not significantly different between the two groups on days 7 and 21. The expression of *POU5F1* mRNA in the TN-C-treated experimental group decreased from day 7 to day 21, but not significantly, and the control group changed only slightly from day 7 to 21 (Fig 3a). The expression of *nestin* mRNA was significantly higher in the TN-C-treated experimental group than in the control group at day 7 (*P* < 0.05). The expression of *nestin* mRNA in both groups showed a significant decrease from day 7 to day 21 (*P* < 0.05) (Fig 3b). The expression of *DSPP* mRNA in both groups increased significantly from day 7 to day 21 (*P* < 0.05) (Fig 3c) but the expression of *DSPP* mRNA was not significantly different between the two groups on day 7 or day 21. The expression of *OCN* mRNA in both groups increased significantly from day 7 to day 21 (*P* < 0.05). The expression of *OCN* mRNA was significantly higher in the control group than in the TN-C-treated experimental group on day 7 and on day 21 (*P* < 0.05) (Fig 3d).

**FIGURE 3.**
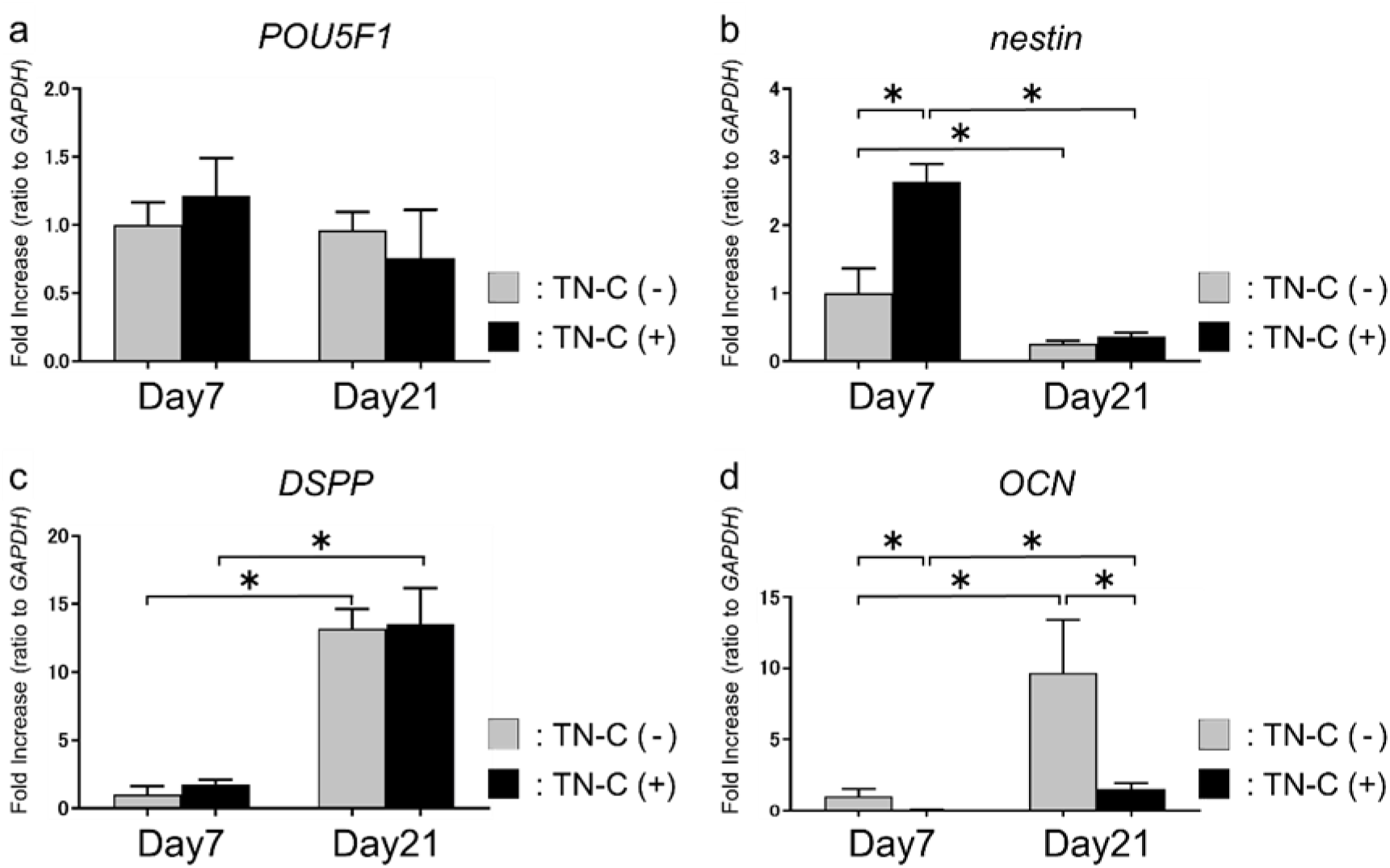
qRT-PCR analysis of DPSCs *in vitro*.

The expression of *POU5F1* mRNA was not significantly different between the two groups at days 7 and 21 (a). The expression of *nestin* mRNA was significantly higher in the TN-C-treated experimental group than in the control group at 7 days (b). The expression of *DSPP* mRNA in both groups increased significantly from day 7 to day 21 (c). The expression of *OCN* mRNA was significantly higher in the control group than in the TN-C-treaed experimental group at day 7 and day 21 (d). Data are expressed as means ± SD; \**P* < 0.05 compared with the control group (a-d).

#### Alizarin Red Staining

After 14 days of culture, mineralized nodules in the control group were observed by Alizarin Red staining (Fig 4a). On the other hand, calcified nodules in the TN-C-treated experimental group were almost not visible at 14 days. After 21 days of culture, calcified nodules in both groups were visible. Furthermore, when the ratio of the area of calcified nodules was calculated, the area of calcified nodules in the control group was significantly greater than in the TN-C-treated experimental group on day 14 and day 21 (*P* < 0.05) (Fig 4b).

**FIGURE 4.**
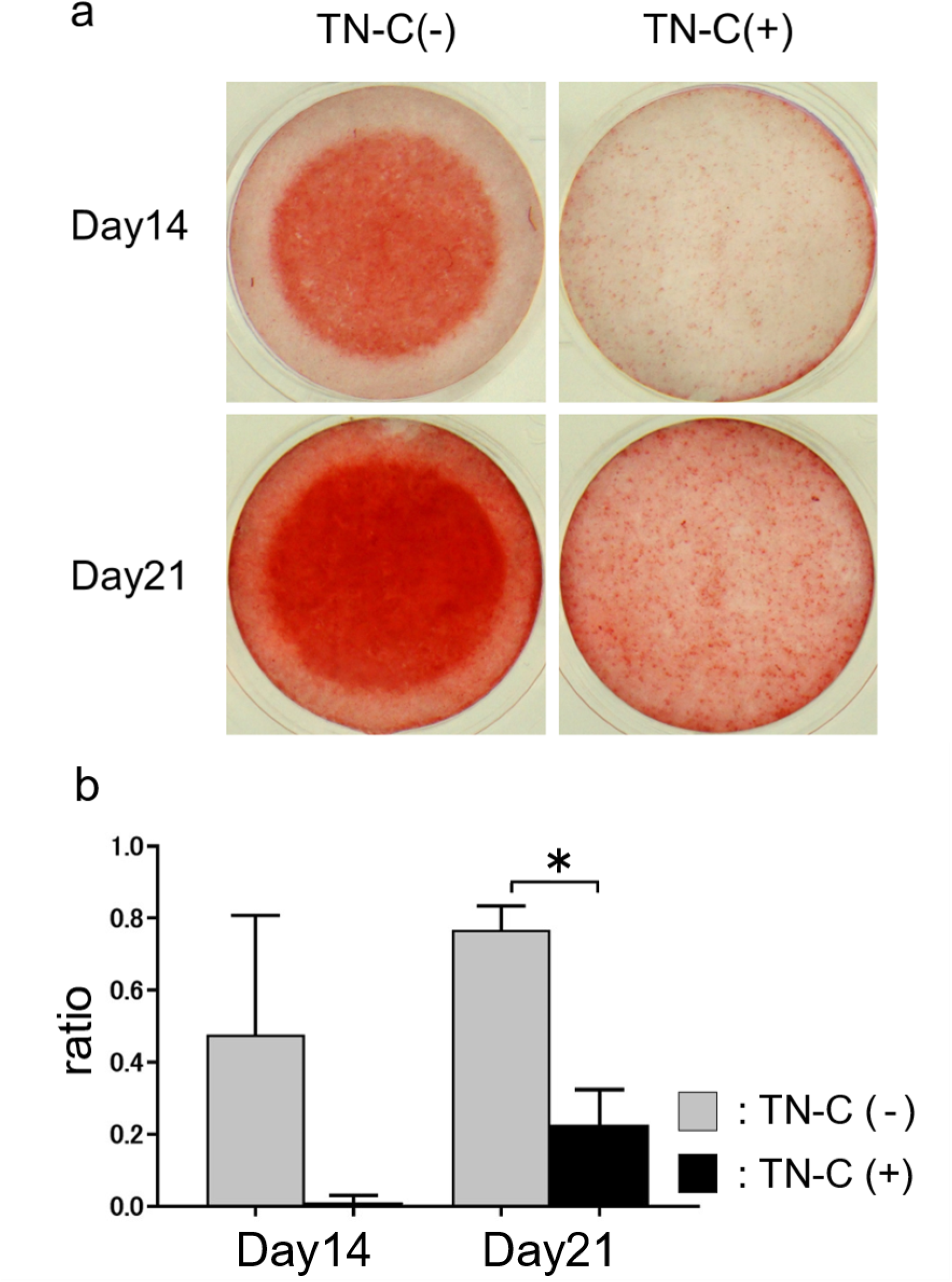
Alizarin red staining.

DPSCs were cultured in odontogenic medium with or without TN-C for 14 and 21 days. Calcium deposition was evaluated using Alizarin red staining (a). After staining, the ratio of the stained area was calculated using the ratio of the area of calcified nodules. The area of calcified nodules in the control group was significantly greater than in the TN-C-treated experimental group on day 21 (b). Data are expressed as means ± SD; \**P* < 0.05. (a) Alizarin red staining. (b) Ratio of the area of calcified nodules.

## DISCUSSION

Granulation tissue in the dental pulp just below the pulp stump and vascular extensions were observed on day 7, and spindle-shaped cells were observed at the root canal opening on day 21 in the TN-C-treated experimental group but no inflammatory cells were observed on day 7 or 21 in the experimental group. By contrast, the pulp tissue in the control group was observed with inflammatory cells on days 7 and 21. The wound healing process involves three stages: the inflammation reaction, cellular element proliferation and synthesis, and remodeling [36]. Treatment that takes advantage of the ability of DPSCs to differentiate into odontoblasts implies an initial inflammatory reaction followed by the proliferation of fibroblasts and DPSCs and their migration to the injury site to replace the lost tissue [37]. TN-C significantly impacts angiogenesis and healing of wounds and counters adverse effects on healing such as excessive inflammation and wound infection [38,39]. Hence, TN-C modulates wound inflammation, suggesting that it may have helped the healing.

Nestin immunoreactivity was observed at day 7 postoperatively and DSPP immunoreactivity was observed at day 21 postoperatively in the dental pulp just below the pulp stump in the TN-C treated experimental group. TN-C has been previously shown to be expressed in the subodontoblastic cell layers during dentin formation in tooth development [31]. Furthermore, it has been reported that TN-C may cause odontoblast differentiation since TN-C is expressed by odontoblasts just below the dentin bridging after direct pulp capping [33]. Matsuoka et al. reported that TN-C promotes differentiation of the cell population isolated from rat dental pulp into mineralized tissue-forming cells *in vitro*, and the results of our previous study suggested that TN-C induces differentiation of the dental pulp into odontoblast-like cells *in vivo* [34].

The cell proliferation rate was not significantly different between the TN-C treated experimental group and the untreated control group. Previous studies have reported that TN-C promotes pulp cell proliferation [34]. However, it has also been reported that TN-C does not affect the proliferation of bone marrow mesenchymal stem cells [28]. TN-C binds to cell membrane receptors, such as integrin, Toll-like receptor 4 and epidermal growth factor receptor and regulates various cell processes including proliferation [40, 41]. However, it has been shown that TN-C does not directly promote cell proliferation, but rather exerts its effects through interactions with growth factors [42]. Dental pulp cells have been found to express growth factors such as BMP4 [43, 44]. Hence, TN-C interacts with growth factors in dental pulp cells to promote proliferation, but not in DPSCs.

The results of this study show that nestin mRNA expression significantly increased in the TN-C-treated experimental group on day 7 compared to the control group. DSPP mRNA expression was also significantly increased from day 7 to day 21 both in the experimental group and in the control group. These results suggest that TN-C promotes the early differentiation of DPSCs into odontoblast-like cells. However, OCN mRNA expression was less significantly increased in the experimental group than in the control group on both days 7 and 21. Furthermore, Alizarin red staining showed the decreased formation of calcified nodules in the experimental group compared with the control group. This suggests that TN-C works to inhibit mineralization. TN-C is known to be expressed in the dental papilla mesenchyme during tooth development [45]. The FNIII repeats of TN-C have been reported to have an affinity for FGF-8 [46, 47]. FGF-8 is expressed during early tooth development and is involved in mesenchymal condensation and the induction of dental papilla cell differentiation [48, 49, 50]. Hence, this suggests that the binding of FGF8 in odontogenic medium to the FNIII domain of TN-C may have promoted the early differentiation of odontoblast-like cells in DPSCs. BMP4 is involved in dentin formation [51] and TN-C has been shown to inhibit BMP4 signaling [52]. Therefore, TN-C inhibited BMP4 in odontogenic medium, suggesting that it may have acted to inhibit mineralization. TN-C is not expressed in mineralized dentin and dentin bridging or in mature odontoblasts but is intensely expressed in the subodontoblastic cell layers. This suggests that TN-C is related to the ability of pulp cells to differentiate into mineralized tissue-forming cells and to participate in the arrangement of mineralized tissues [31, 33] and the results of this support that possibility.

## CONCLUSION

In conclusion, the data of this study show that TN-C regulates inflammation during the healing process, induces the early differentiation of DPSCs to odontoblasts, and may further contribute to the inhibition of excessive dentin formation.

## DATA AVAILABILITY STATEMENT

The datasets generated and/or analyzed during the current study are available from the corresponding author on reasonable request.

